# Influence of Cognitive Demand and auditory noise on Postural Dynamics

**DOI:** 10.1101/2024.10.08.617302

**Authors:** Sam Carey, Ramesh Balasubramaniam

## Abstract

The control of human balance involves an interaction between the human motor, cognitive, and sensory systems. The dynamics of this interaction are yet to be fully understood, however, work has shown the performance of cognitive tasks to have a hampering effect on motor performance, while additive sensory noise to have a beneficial effect. The current study aims to examine whether postural control will be impacted by a concurrent working memory task, and similarly, if additive noise can counteract the expected negative influence of the added cognitive demand. Postural sway of healthy young adults was collected during the performance of a modified N-back task with varying difficulty, in the presence and absence of auditory noise. Our results show a reduction in postural stability scaled to the difficulty of the cognitive task, but this effect is less prominent in the presence of additive noise. Additionally, by separating postural sway into different frequency bands, typically used to assess the exploratory vs feedback-driven stabilizing dynamics of sway, we found a differential effect between the cognitive task and additive noise, thus demonstrating that both frequency regimes of postural sway are sensitive to high cognitive load and increased sensory information.

## INTRODUCTION

The ability to stand upright for extended periods of time is often taken for granted. With the majority of the human body’s mass located in the head, arms, and trunk, the human body is inherently unstable, often analogous to an inverted pendulum. The goal of the postural control system is to maintain stability within this variable paradigm as the surrounding environment shifts and changes. This results in two significant dynamics of the postural control system: static and dynamic equilibrium. Static equilibrium involves maintaining a stable position under the influence of gravity alone while dynamic equilibrium concerns the body’s segmental adjustments in response to external stimuli (e.g., uneven surfaces, changing sensory input, cognitive load, etc.) (Lalonde and Stazielle 2007).

Postural stability constitutes an essential element within the human body’s motor control and coordination system. It is imperative for the execution of both static and dynamic activities (Wikstrom et al. 2005) and is supported by two distinct levels of processing: higher “controlled” processing (Raftopoulos 2005; Boisgontier et al. 2013), involving the basal ganglia-cortical loop (Jacobs and Horak 2007), and lower “automatic” processing, underpinned by brainstem synergies (Honeycutt et al. 2009). However, the motor synergies that govern basic coordination patterns, such as standing and walking, are considered autonomous organizations (Morasso et al. 2019), not requiring conscious effort or cognitive demand. Which leads us to the question at hand, if these basic coordination patterns that govern upright stability are considered subcortical/supraspinal, how do they interact with subsequent motor tasks, cognitive demands, or alterations in the environment? Before we attempt to answer this question, it would serve us well to clarify some of the more nuanced dynamics of postural control.

### What is postural control and why is it important?

Maintaining postural control and stability is essential for task success in most of our daily lives. Postural stabilization is not an end in itself; rather, it is valuable to the extent that it facilitates the achievement of countless goals throughout the human lifespan (Riccio and Stoffregen 1988). Although it may appear simple, postural stability is more complex than many realize.

Falling under the purview of the postural control system, stability relies on the integration of sensory information and cognitive processing of the task at hand (Duarte and Frietas 2010). Through the processing of this information, the postural control system is able to effectively maintain the position of the center of mass (CoM) through the control of the center of pressure (CoP) and thus preserve stability during upright standing and task completion. An active process that humans have learned to automatize relatively effectively (unless a perturbation were to arise). However, as stated previously, automaticity does not imply simplicity, the postural control system remains a highly complex system comprised of multiple interacting elements required for success.

While posture is inherently unstable, human stability is rarely threatened under quiet or static conditions. Yet, human stability becomes threatened when a secondary task is performed during upright standing, whether that task be cognitive or motor related. Performing a cognitive task while standing is common in everyday life, such as reading a list at a grocery store or recalling and sharing information with a coworker in the breakroom. Although these may seem like simple tasks, their execution during standing alters the performance of both the cognitive task and the stability of standing.

### Posture and Cognition

Historically, researchers have focused on the superordinate organization of tasks concerning postural control (Stoffregen et al. 1999), often neglecting the influence that postural control may exert on concurrent tasks and vice versa. Typically, supra-postural mechanisms have been studied without considering the changes in postural dynamics that may occur during the performance of secondary tasks (Stoffregen et al. 2000). Given that falls are a significant cause of injury-related deaths among older adults (Hornbrook et al. 1994; CDC 2024), understanding the interaction between the postural system and the performance of concurrent tasks is of paramount importance. The motor synergies responsible for basic coordination patterns, such as standing and walking, have been studied as autonomous organizations that underpin balance, safety, and locomotion (Morasso et al. 2019). However, past research has often overlooked the coexistence of these synergies with perceptual-cognitive functions beyond balance and movement, which are crucial in facilitating suprapostural functions (Stoffregen et al. 1999, 2000).

Research indicates that maintaining posture against external perturbations requires the same cognitive processes for mentally demanding tasks (Quant et al. 2004, 2005). Behavioral studies examining the modulating effects of postural and cognitive activities simultaneously provide evidence that postural control and cognitive tasks share common resource requirements (Fraizer and Mitra 2008). Additionally, multiple studies have demonstrated a possible interaction effect between the performance of postural control and cognitive tasks (Wollacott and Shumway-Cook 2002). This interaction may seem surprising, as posture and balance are typically considered spinal or subcortical processes, while cognition is viewed as a cortical process.

However, this distinction is not entirely accurate, as the cerebellum plays a role in both sensory processing and cognition (Morton and Bastian 2004) and cortical involvement has been shown in postural reflexes and adaptation (Johansson et al. 1994; Raftopoulos 2005; Boisgontier et al. 2013).

The performance of concurrent cognitive tasks has been shown to interact with postural control, typically in an unfavorable way. Maylor and Wing (1996) previously found that older adults were more strongly affected by cognitive task performance than younger adults. In a digit recall task involving spatial memory, they observed increased sway in older adults compared to conditions without a cognitive task when compared to younger populations. However, the nature of this relationship and the influence cognition has on postural control remains unclear.

Continued work has attempted to better understand this interaction more thoroughly through specifying the task constraints: verbal vs. visual stimuli (Ramenzoni et al. 2007), comparing age groups (Maylor and Wing 1996), encoding vs. retrieval phases of tasks (Maylor and Wing 2001), and assessing clinical populations (Pellecchia 2003; Redfern et al. 2001).While this body of work was able to show this interaction effect, the understanding of why it occurs still remains unknown.

Multiple theories have attempted to explain the interference effect seen between posture and a concurrent cognitive task. A common theory is the ‘posture first’ principle which suggests that individuals prioritize maintaining balance when performing a cognitive task, potentially leading to decreased cognitive performance (Andersson et al. 2002). However, this theory presupposes that reduced postural sway amplitude correlates with increased stability, a notion challenged by Stoffregen et al. (1999), who argued that postural control serves as an adaptive or prepatory behavior rather than an end in itself. Suggesting that a decrease in sway may not directly imply an increase in stability, but a more general alteration in the sway dynamics.

Stability is more complex than an increase or decrease in the amplitude of sway. Nevertheless, it is assumed that cognition has a negative effect on postural sway dynamics, and there is a need to discover why this interaction occurs, and possibly how to counteract it. Similarly, limited attentional availability theories have posited that the mind only has enough resources to properly perform a single task at a time, and the more tasks performed, the higher degree of decrement is seen in each of the tasks. Today, the interaction effect is still unknown, with multiple theories being used to explain this effect to no avail.

### Posture and Noise

Successful postural control relies on sensory feedback and predictions from somatosensory, vestibular, visual, and auditory modalities (Dozza et al. 2007). Even with ample sensory information, postural sway remains sensitive to feedback from any of these modalities (Yeh et al. 2010). Increased sensory information availability can reduce the multidirectional variability of sway, thereby improving balance and enhancing adaptability to potential perturbations (Carey et al. 2023, 2024; Ross et al. 2015, 2016a, b). The multisensory nature of postural control necessitates system adaptability; as environmental conditions change, so does the reliance on different sensory systems. For instance, when visual information is limited (e.g., at dusk or night), individuals rely more on tactile, vestibular, or auditory inputs to maintain stability and navigation.

Moreover, introducing noise stimulation, particularly in the auditory or tactile modalities, can enhance the stability of these systems. For example, sub-sensory mechanical noise applied to the soles of the feet has been shown to reduce postural sway in healthy young adults (Priplata et al. 2002), healthy older adults, and individuals with central and peripheral sensorimotor deficits (Priplata et al. 2003, 2006). Recent studies have demonstrated strong stabilizing effects of auditory noise on postural sway variability (Ross and Balasubramaniam 2015; Ross et al. 2016a, b; Carey et al. 2023, 2024), although earlier research yielded mixed results (Hegeman et al. 2005). Further investigation suggests that the acoustic properties of auditory stimuli may be more critical in reducing sway than whether the signal provides velocity or position information (Hegeman et al. 2005; Dozza et al. 2007), which may explain the previously mixed findings. For example, Deviterne et al. (2005) observed reduced sway when participants listened to prolonged speech, but not when listening to a single sustained tone.

Previous literature has established that the postural control system depends on sensory information, cognitive demands of the environment, and the motor capabilities of the individual. Cognitive tasks can decrease postural stability, while sensory noise can enhance it (Ross et al. 2015; Carey et al. 2024). This presents an opportunity to use the beneficial effects of additive sensory noise to counteract the negative impact of cognitive demands on stability. This study aims to evaluate the effect of a modified n-back cognitive task on postural stability while also employing additive auditory noise to mitigate the cognitive interference. We hypothesize that while cognitive performance will reduce stability, the addition of noise will restore stability to levels comparable to those observed in quiet conditions.

## METHODS

### Participants

Twenty-eight healthy young adults (mean age = 23.18 ± 4.27) of varying heights (66.50 ± 3.69 inches) and weights (152.93 ± 28.31 lbs.) were recruited from the University of California, Merced student population. Self-reported screeners were used to exclude participants with hearing impairments, arthritis, orthopedic conditions, or neurological disorders (Carey et al. 2023, 2024; Ross and Balasubramaniam 2015; Ross et al. 2016a, b). No participants reported recent injuries or skeletomuscular disorders, and all could stand unassisted for the duration of the experiment. The experimental protocol was carried out in accordance with the Declaration of Helsinki, reviewed by the UC Merced IRB, and all participants gave informed and written consent prior to testing.

### Protocol

Participants were instructed to stand on a force platform in a relaxed, comfortable standing position with their arms at their sides and feet shoulder width apart while wearing headphones. Participants were given breaks periodically throughout the task, typically after 5-7 trials to sit down and rest their legs. Upon the continuation of the next block of trials, participants were instructed again to place their feet shoulder width apart and place their arms back at their sides. Participants performed a total of 30 trials, 10 trials of each condition (No Cognitive, Easy Cognitive, & Hard Cognitive). The easy and hard conditions were blocked with 10 trials each, the No Task was broken into 3 blocks, with 4 trials at the beginning and end, and 2 trials in the middle of the two easy and hard conditions. This spread was designed to gather a baseline, while also controlling for any possible fatigue the participants may experience throughout the task. The easy and hard blocks were randomized between participants.

Amongst all the task conditions, noise was randomly played during half of the trials. Trials lasted 90 s and were either accompanied by white noise (intensity of 75 dB) or silence. The noise and silence conditions were presented in a randomized order with a total of 15 noise trials and 15 silent trials. Participants were exposed to the noise stimuli prior to the experiment to verify that the stimuli were not uncomfortable. No participants reported discomfort at these intensities. Center of Pressure (CoP) was sampled at 200 Hz with an AMTI Force and Motion platform (Optima BP400600-2000 GEN 5; AMTI Force & Motion, Watertown, MA, USA). All data were collected in a single session. The auditory noise stimulus was generated using MATLAB to be random signals with a constant spectral density.

### Task

While standing, participants performed a modified N-back task of two difficulty levels:

Easy and Hard. Each trial lasted 90 seconds with a total of 10 trials of each difficulty level of the task. The task was presented on a projector screen 210 cm in front of the participants, with the middle of the screen being adjusted for each participant to be at eye height. No participants reported difficulty reading or seeing the letters clearly during the task.

Capitalized letters appeared on the screen in a random order, with a fixation cross appearing between each letter. The letters appeared on the screen for 0.5 s while the fixation cross lasting for 2.0 s, and this pattern repeated for a total of 90 s, alternating between the letters and fixation cross. Each trial started and ended with a fixation cross, a total of 34 letters were presented during each trial and 35 fixation crosses. The letter “X” was excluded from the task presentation due to its similarity to the shape of the fixation cross “+,” all other letters within the alphabet were used.

The goal of the **EASY** condition was to count the number of times any letter was repeated after a single fixation cross. For example, if the following sequence of letters were to appear on the screen in the order of: *A + A + F + K + R + R + T + P* etc., there would be a total count of 2, one for the letter A being repeated and one for the letter R being repeated. At the end of each trial, the participants would self-report the count of repeated letters they saw. For the **HARD** conditions, the same presentation format was used, but now it represented an N-2 type task where the repetitions no longer counted following a single fixation cross, but with a single random letter between the repetitions and two fixation crosses. For example, if the following sequence was presented: *A + R + A + K + T + R + T + P + F + P*, etc., the count would be 3, for the *A, T*, and *P* being repeated.

At the end of each session, the absolute error count was calculated as the difference between the expected value and the self-reported value across all trials for the easy and hard conditions. Participants were instructed to not utilize their hands or fingers to keep the count of repetitions during the trial to account for possible offloading of the cognitive task.

### Analyses

All CoP was analyzed using custom scripts in MATLAB (MathWorks, Natick, MA, USA). The first 4 s of each trial were removed to eliminate any potential startle response the participants might have had to stimulus onset. Radial sway (RS) of the CoP was calculated for each sample (i) using the anterior–posterior (A–P; x) and medial–lateral (M–L; y) components of sway following (Lafond et al. 2004):

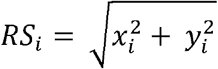

Average RS was calculated for each trial and was used to assess bidirectional variability in CoP during trials (Lafond et al. 2004). There are multiple other measures of postural stability that are efficient and effective when studying postural sway, including mean velocity, median power–frequency, RMS distance and sway area (Lin et al. 2008). While RS is not a direct metric of stability, it utilizes the multidirectional variability of sway to offer a more robust understanding of the sway dynamics that may lead to stability, compared to a unidirectional metric like the standard deviation of CoP magnitude or velocity (Lafond et al. 2004). Trial outliers were determined as trials with trial averages of ±2 standard deviations from that subject’s mean within condition and were removed. We removed an average of X% of the total trials (X out of the total X trials). No subject had more than 2 trials (out of 10) removed per condition.

The statistical analysis was then repeated using the filtered high and low frequency RS separately to assess changes in slower and faster timescales of postural control (following the methods of Yeh et al. 2010, 2014). We used low- and high-pass Butterworth filtering routines, as in Yeh et al. 2014, to decompose sway into low (0.3 Hz) -frequency sway. The filter cutoff was chosen based on van den Heuvel et al. (2009) to separate into sensory feedback-related sway and spontaneous/exploratory sway.

Finally, detrended fluctuation analysis (DFA) was used to quantify the sway dynamics over time (Delignières et al. 2003; Collins and Luca 1994). DFA is used to study the behavior of the timeseries of CoP. This method, first introduced by Peng et al. (1994), is a scaling analysis method that provides a scaling exponent, which offers information concerning the correlational properties of the CoP signal. When the DFA value exists between 1 < α <1.5, the postural sway is considered antipersistent. This means that the sway moves in successive steps in random directions (a semi-random walk) and does not trend toward the same direction. Antipersistent radial sway dynamics is commonly described in healthy postural sway. This analysis was completed as in (Blázquez et al. 2010) using the same parameters. See Blázquez et al. (2010) and Delignières et al. (2003) for more details on the DFA method.

## RESULTS

To examine the effects of Noise Stimuli and Cognitive Condition on Radial sway, while accounting for individual subject differences, a linear mixed-effects model was employed using the *lmer* function in R. the model specified Radial Sway as the dependent variable, with the Noise Stimuli and Cognitive Condition as fixed effects, including their interactions. Subjects were treated as a random effect to account for repeated measures and inter-subject variability. The formula used in R was *RS ∼ Stim*Condition + (1*|*Subject)*. This approach allows for the investigation of both the main effects and interaction effects of Stimuli and Condition on Radial Sway, while controlling for potential confounding influences of individual differences across subjects.

### Radial Sway

The fixed effects of the model revealed several significant findings. The intercept was estimated at 5.726 (SE = 0.357, t = 16.044, p <0.001), indicating the baseline level of Radial Sway. The main effect of noise was significant, with additive noise resulting in a decrease in RS (Estimate = -1.273, SE = 0.203, t = -6.281, p <0.001). For the Cognitive task, the Easy level did not significantly differ from the baseline (Estimate = 0.144, SE = 0.199, t = 3.786, p = 0.486) (Figure 3), however, Hard condition was significantly different, with an increase in RS when the hard task was performed (Estimate = 0.7532, SE = 0.1980, t = 3.786, p < .0001). The interaction between Noise and Cognitive task was also examined. There was a significant interaction between noise and the *Easy* task (Estimate = 0.831, SE = 0.282, t = 2.945, p = 0.003), suggesting a combined effect of these conditions on RS. However, there was no interaction effect between noise and the *Hard* task (Estimate = 0.458, SE = 0.282, t = 1.624, p = 0.105).

**Figure 2.**
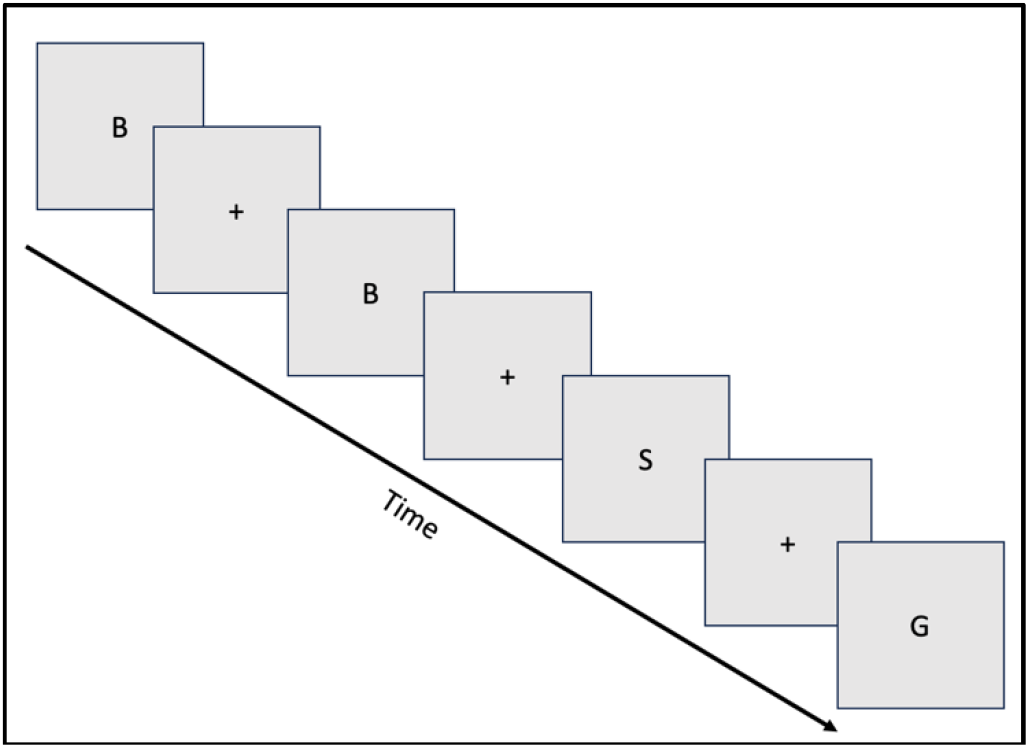
Example format of the presentation of the modified n-back task.

**Figure 3.**
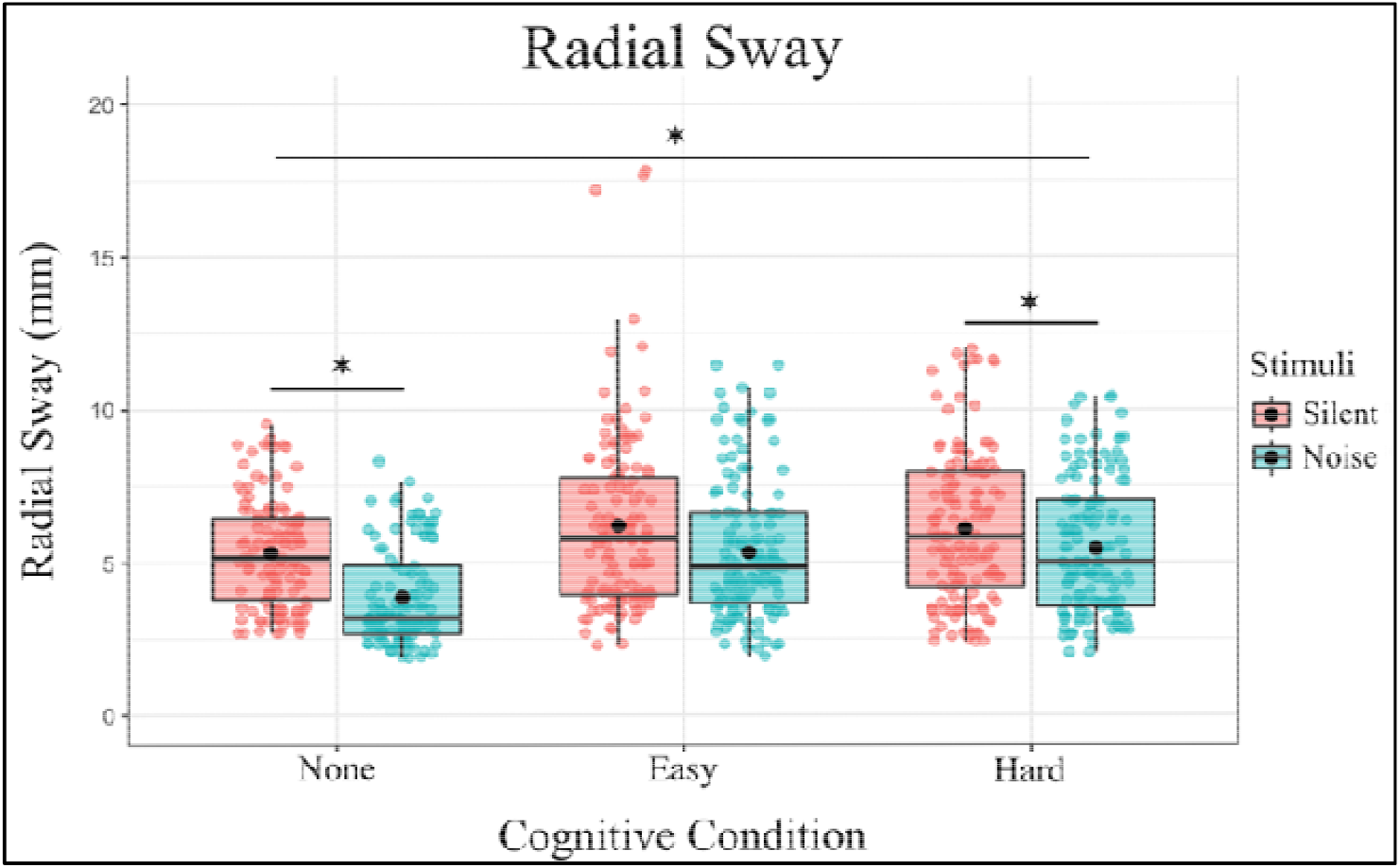
Radial Sway is significantly reduced with the introduction of auditory noise in the No-Cognitive and Hard-Cognitive Conditions and increased with the introduction of a Hard Cognitive task. Box and whiskers plot with the solid black line representing the median, the solid black dot representing the mean, and the extending lines showing the maximum and minimum values.

In summary, the model indicates that both Stimulus and Condition significantly influence Radial Sway, with notable interaction effects. The inclusion of random effects for subject’s accounts for individual variability, enhancing the robustness of the findings.

To further investigate the effects of different levels of Noise and Cognitive Condition on RS, post hoc comparisons were performed. These comparisons compared various combinations of Noise and Cognitive Condition using Tukey’s method for multiple comparisons with degrees of freedom calculated using the Kenward-Roger method (Table 1). The following significant findings were observed:

**Table 1.**
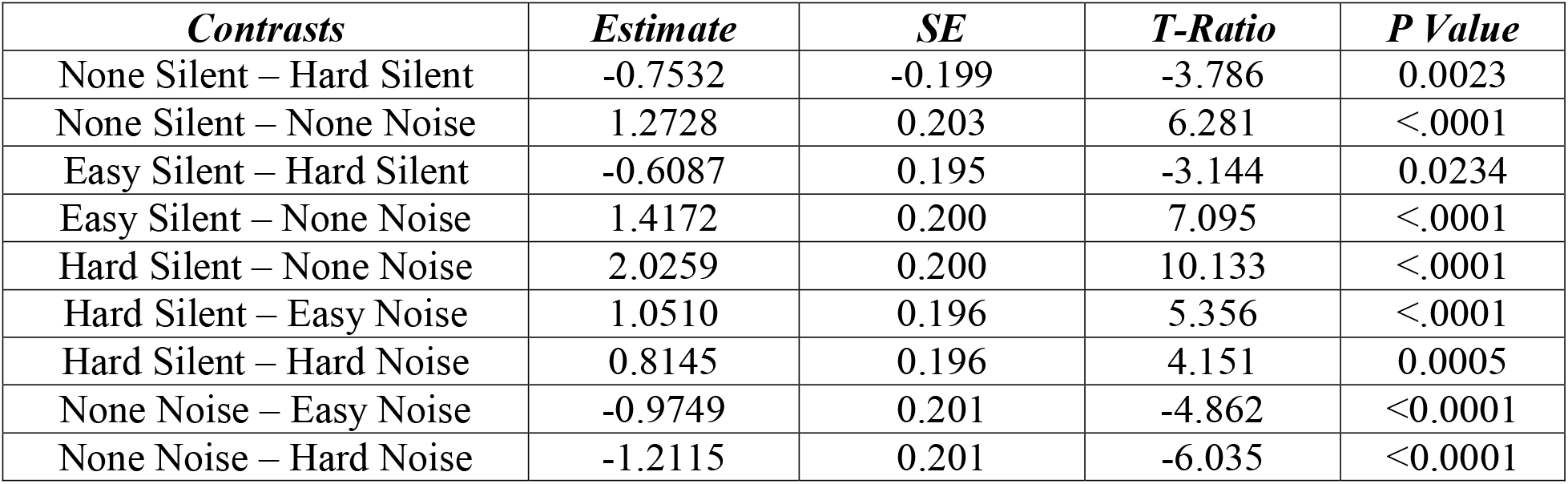

These results indicate that both Stimulus and Condition significantly influence RS, with notable differences between specific levels of these factors. The use of Tukey’s method for multiple comparisons ensures that the reported p-values are adjusted for the family-wise error rate, providing robust statistical inference.

### High-Frequency Radial Sway

The fixed effects of the model revealed several significant findings. The intercept was estimated at 2.6099 (SE = 0.1502, t = 17.367), representing the baseline level of High-Frequency RS when all predictors are at their reference levels. The additive noise had a significant effect on High-Frequency RS (Estimate = -0.3472, SE = 0.0767, t = -4.527, p <0.001), indicating that the presence of additive noise substantially decreases High-Frequency RS. Similarly, The Hard cognitive condition was associated with a significant increase in High-Frequency RS (Estimate = 0.3232, SE = 0.0760, t = 4.251, p < 0.001). In contrast, the Easy condition did not show a significant effect on High-Frequency RS (Estimate = -0.0182, SE = 0.0755, t = -0.242, p = 0.809). These results suggest that while the Easy condition does not alter High-Frequency RS,, the Hard condition leads to an increase in High-Frequency RS dynamics compared to baseline (Figure 4), indicating a differential effect of cognitive load on postural sway dynamics.

**Figure 4.**
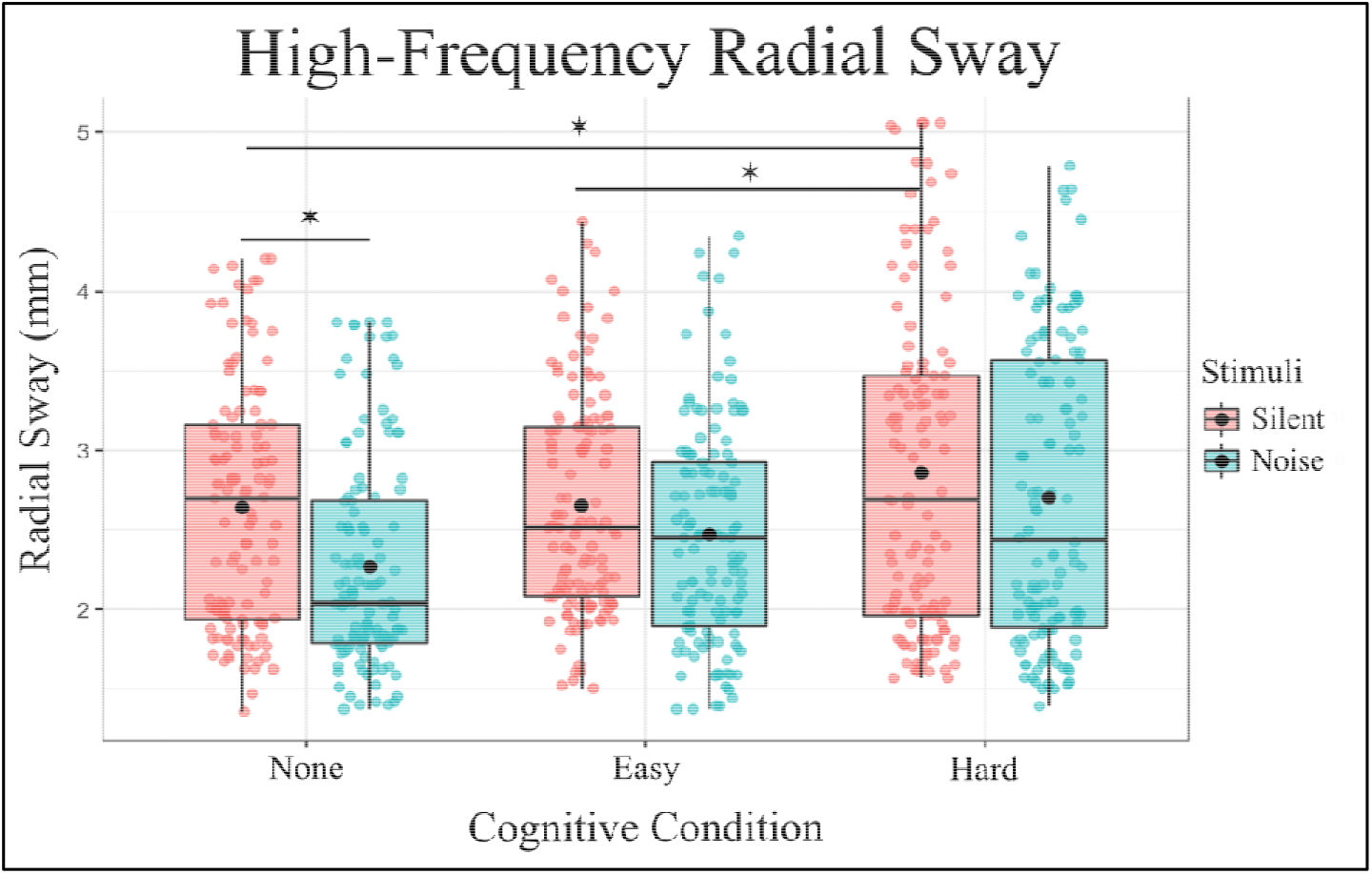
High-Frequency Radial Sway is significantly reduced with the introduction of auditory noise and increased with the introduction of a Hard Cognitive task. Box and whiskers plot with the solid black line representing the median, the solid black dot representing the mean, and the extending lines showing the maximum and minimum values.

The interaction effect revealed that the interaction between additive noise and the Easy condition trended toward significance (Estimate = 0.1900, SE = 0.1068, t = 1.863, p = 0.063), suggesting a potential moderating effect of the Easy condition on the impact of noise on High-Frequency RS. However, the interaction between Noise and the Hard condition was not significant (Estimate = 0.1575, SE = 0.1071, t = 1.472, p = 0.142), indicating that the hard condition does not influence the effect of noise on High-Frequency RS.

In summary, the analysis indicates that noise significantly reduces High-Frequency RS, particularly in the Hard condition, which shows an increase in High-Frequency RS compared to the baseline. The Easy condition does not significantly alter High-Frequency RS, and its interaction with noise is marginally significant. These findings contribute to our understanding of sensory processing and balance control under different environmental conditions, highlighting the differential impacts of noise and task difficulty on postural sway.

To further investigate the effects of different levels of Noise and Cognitive Condition on RS, post hoc comparisons were performed. These comparisons compared various combinations of Noise and Cognitive Condition using Tukey’s method for multiple comparisons with degrees of freedom calculated using the Kenward-Roger method (Table 2). The following significant findings were observed:

**Table 2.**
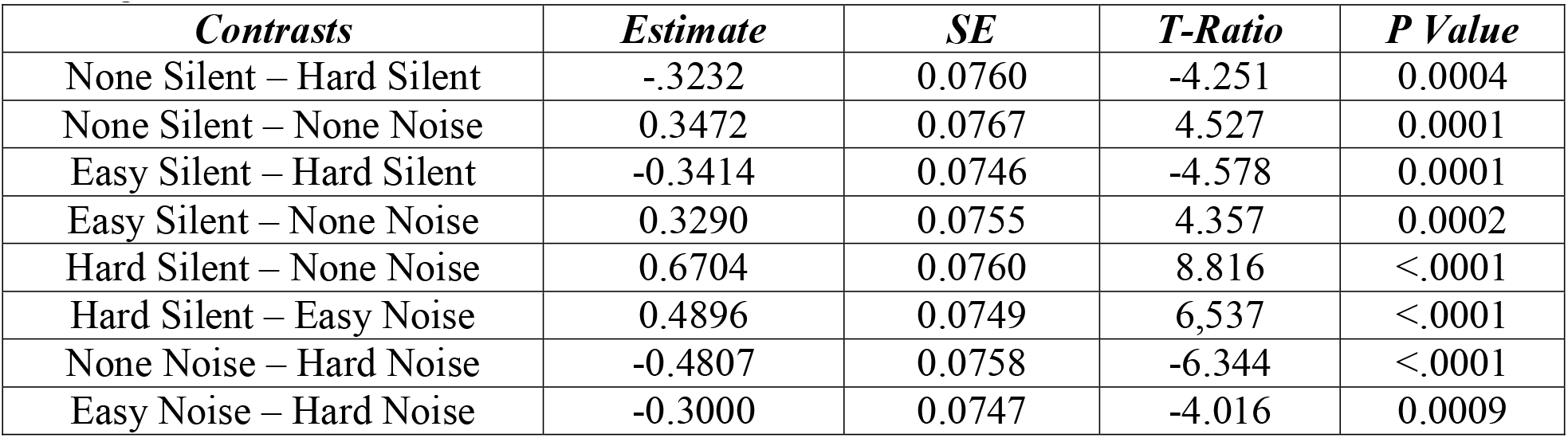
Post-Hoc Comparisons of significance. For the entirety of the post-hoc comparisons for High-Frequency RS, see Appendix.

| *Contrasts* | *Estimate* | *SE* | *T-Ratio* | *P Value* |
| --- | --- | --- | --- | --- |
| None Silent – Hard Silent | -.3232 | 0.0760 | -4.251 | 0.0004 |
| None Silent – None Noise | 0.3472 | 0.0767 | 4.527 | 0.0001 |
| Easy Silent – Hard Silent | -0.3414 | 0.0746 | -4.578 | 0.0001 |
| Easy Silent – None Noise | 0.3290 | 0.0755 | 4.357 | 0.0002 |
| Hard Silent – None Noise | 0.6704 | 0.0760 | 8.816 | <.0001 |
| Hard Silent – Easy Noise | 0.4896 | 0.0749 | 6.537 | <.0001 |
| None Noise – Hard Noise | -0.4807 | 0.0758 | -6.344 | <.0001 |
| Easy Noise – Hard Noise | -0.3000 | 0.0747 | -4.016 | 0.0009 |

### Low-Frequency Radial Sway

The fixed effects of the model revealed several significant findings (Fig. 5). The intercept was estimated at 4.330 (SE = 0.252, t = 17.182, p < 0.001), indicating the baseline level of Low-frequency Radial Sway. The main effect of noise was significant, with the introduction of noise resulting in a decrease in Radial Sway (Estimate = -1.073, SE = 0.184, t = -5.826, p < 0.001). For Cognitive Condition, the Easy level did not significantly differ from the quiet baseline condition (Estimate = -0.087, SE = 0.181, t = -0.479, p = 0.632), while the Hard condition was associated with a significant increase in RS (Estimate = 0.524, SE = 0.180, t = 2.903, p = 0.004). Similarly, the interaction between Noise and Cognitive Condition was examined. There was a significant interaction between Noise and the Easy Cognitive (Estimate = 0.655, SE = 0.257, t = 2.549, p = 0.011), but no interaction effect between Noise and the Hard Cognitive Condition (Estimate = 0.200, SE = 0.256, t = 0.782, p = 0.434).

**Figure 5.**
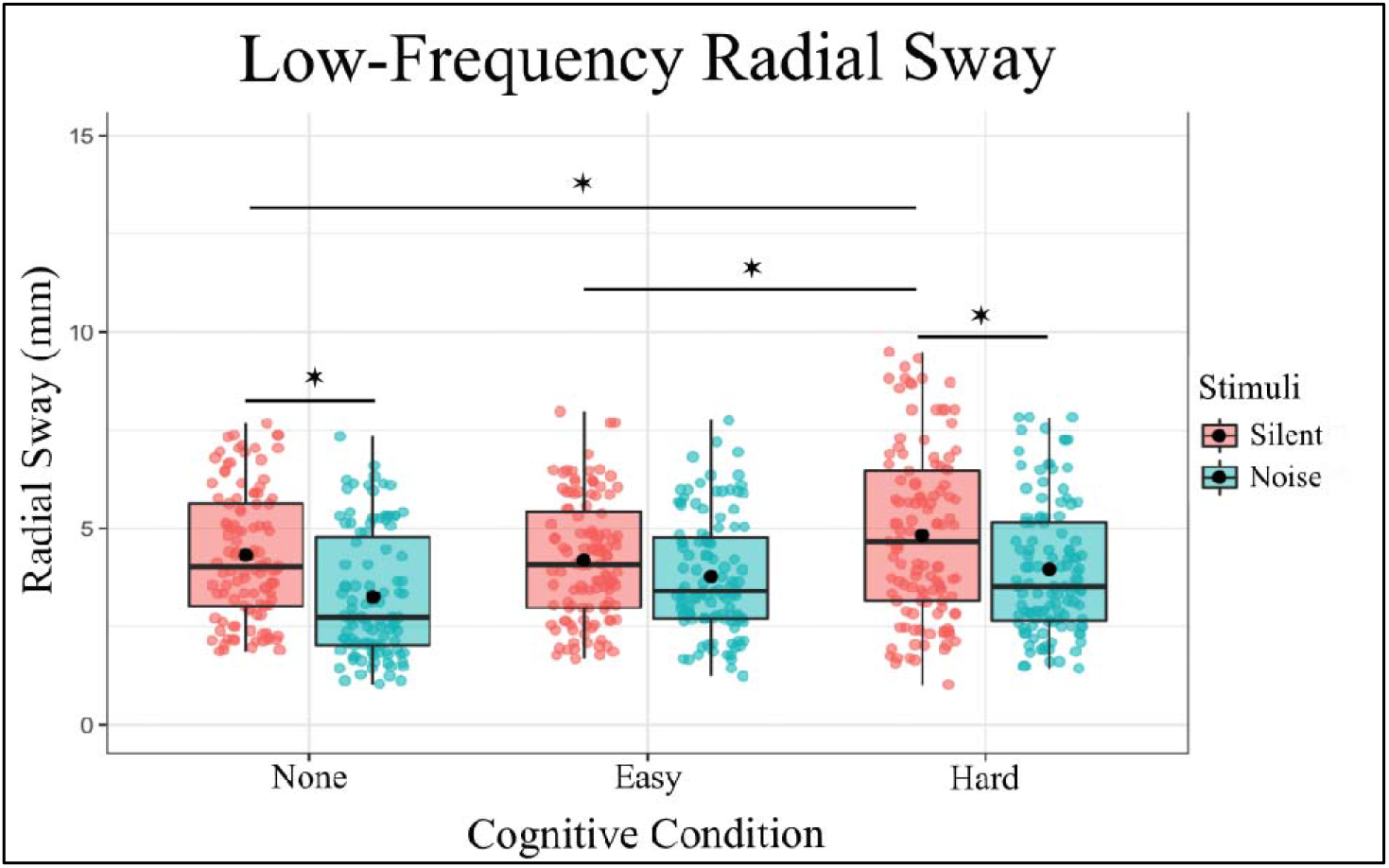
Low-Frequency Radial Sway is significantly reduced with the introduction of auditory noise in the No-Cognitive and Hard-Cognitive Conditions and increased with the introduction of a Hard Cognitive task. Box and whiskers plot with the solid black line representing the median, the solid black dot representing the mean, and the extending lines showing the maximum and minimum values.

In summary, the model indicates that both Stimulus and Cognitive Condition significantly influence RS, with notable interaction effects, The inclusion of random effects for subject’s accounts for individual variability, enhancing the robustness of the findings (Figure 5).

To further investigate the effects of different levels of Noise and Cognitive Condition on RS, post hoc comparisons were performed. These comparisons compared various combinations of Noise and Cognitive Condition using Tukey’s method for multiple comparisons with degrees of freedom calculated using the Kenward-Roger method (Table 3). The following significant findings were observed:

**Table 3.**
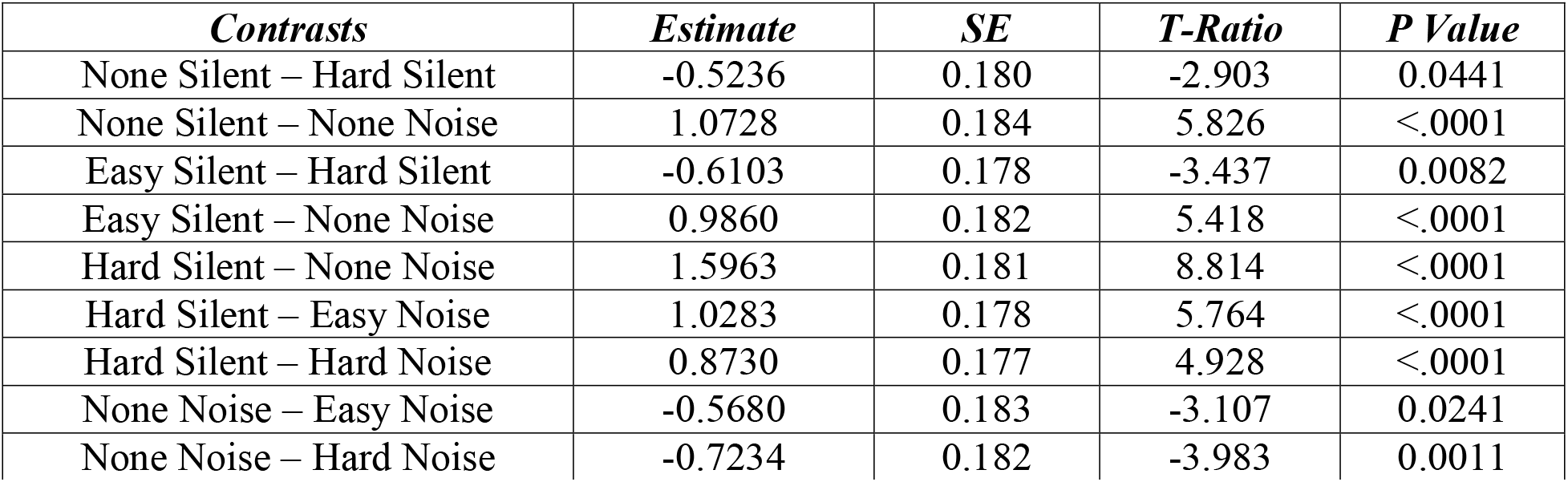
Significant post-hoc comparisons for Low-Frequency RS, for the full set of post-hoc comparisons, see Appendix. The use of Tukey’s method for multiple comparisons ensures that the reported p-values are adjusted for the family-wise error rate, providing robust statistical inference.

### Detrended Fluctuation Analysis

Detrended Fluctuation Analysis showed that RS exhibits anti-persistent fractional Brownian motion (fαm, 1<α <1.5). This semi-random walk pattern is characteristic of postural sway (Blázquez et al. 2010, Delignières et al. 2003, Collins and De Luca 1994). Within this 1– 1.5 range, there are differences between subjects in α. The fixed effects of the model revealed several significant findings for Detrended Fluctuation Analysis (Figure 6). The intercept was estimated at 1.293 (SE = 0.017, t = 76.207, p <0.001), indicating the baseline level of Alpha. The main effect of Noise was significant with additive noise resulting in a decrease in Alpha (Estimate = -0.098, SE = 0.012, t = -8.363, p <0.001). For the cognitive task, the Easy difficulty level did not significantly differ from quiet standing (Estimate = -0.020, SE = 0.011, t = -1.695, p = 0.090), however, the Hard condition did significantly differ from baseline standing (Estimate = -0.031, SE = 0.012, t = -2.687, p = 0.007).

**Figure 6.**
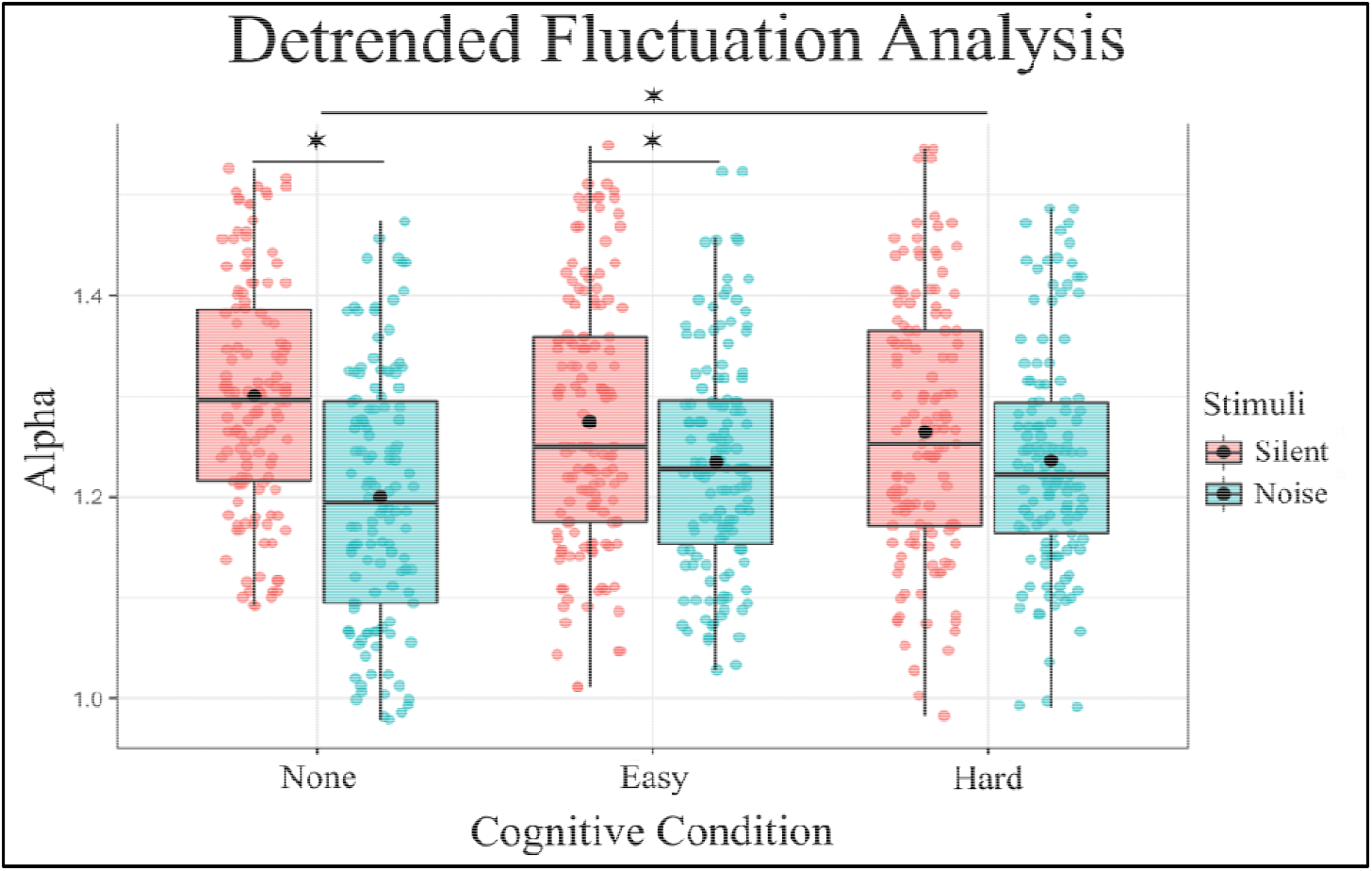
DFA Alpha is significantly reduced with the introduction of auditory noise in the No-Cognitive and Hard-Cognitive Conditions and increased with the introduction of a Hard Cognitive task. Box and whiskers plot with the solid black line representing the median, the solid black dot representing the mean, and the extending lines showing the maximum and minimum values.

There was also an interaction effect between Noise and Cognitive task. Both Noise and the Easy Condition (Estimate = 0.063, SE = 0.016, t = 3.824, p <0.001) had a significant interaction, as did the additive Noise and the Hard Condition (Estimate = 0.073, SE = 0.016, t = 4.422, p < 0.001), suggesting that these combinations significantly affect Alpha levels during standing. In summary, the model indicates that the Stimulus “Noise” significantly decreases Alpha, and this effect is further influenced by the Condition, the inclusion of random effects for subject’s accounts for individual variability, enhancing the robustness of the findings.

To further investigate the effects of different levels of Noise and Cognitive Condition on RS, post hoc comparisons were performed. These comparisons compared various combinations of Noise and Cognitive Condition using Tukey’s method for multiple comparisons with degrees of freedom calculated using the Kenward-Roger method (Table 4). The following significant findings were observed:

**Table 4.**
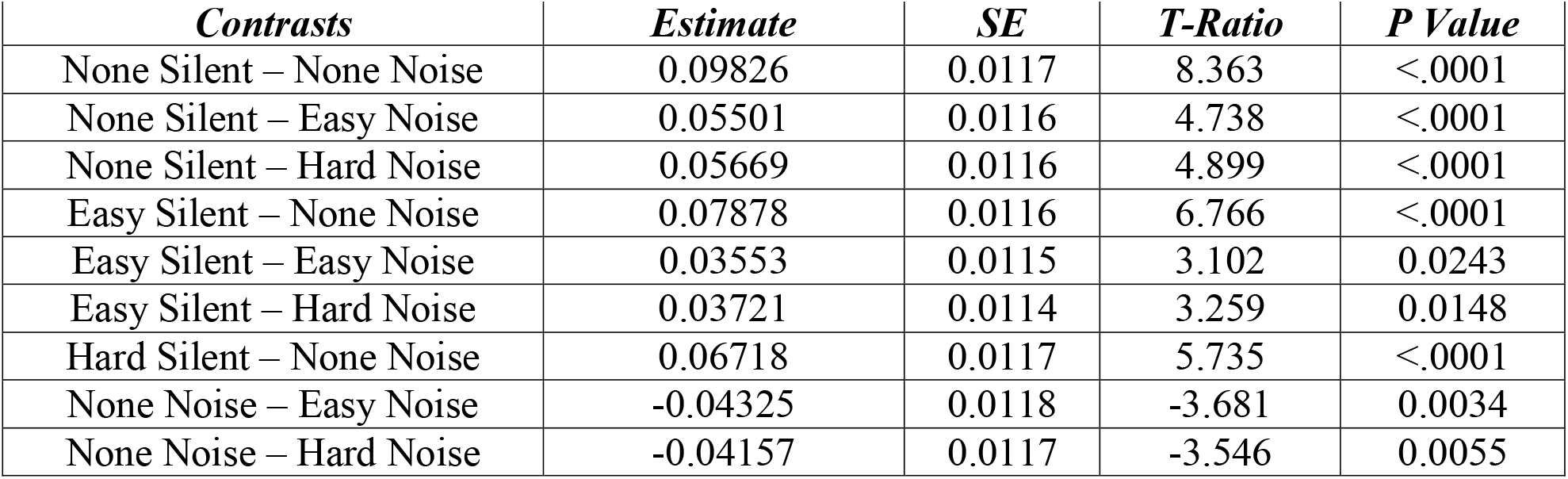
Significant post-hoc comparisons for DFA of RS, for entire post-hoc analysis, see Appendix.

## DISCUSSION

We show a reduction in the multi-directional postural sway variability with the addition of auditory white noise in healthy young adults when compared to silence, further solidifying the beneficial effect noise can have on postural stability (Carey et al. 2023, 2024; Ross et al. 2014, 2016). Furthermore, we found an increase in postural sway variability with the addition of a 90 second working memory task. However, this increase in sway was only found in the Hard condition, not in the Easy condition, implying that the degree of cognitive load differentially effects postural dynamics during dual-tasking performance.

The low and high frequency components of sway were similarly influenced by these added sensory inputs and cognitive demands. Slower timescales of sway are thought to reflect drift of the inertial mass of the body (Winter et al. 1998) and are more susceptible to changes in sensory feedback (Yeh et al. 2010, 2014; van den Heuvel et al. 2009). While faster timescales are interpreted as smaller adjustments around the center of mass that are more directly related to join rigidity and muscle activation (Kiemel et al. 2005; van den Heuvel et al. 2009). Our results show that the introduction of a hard cognitive task increases the variability of the high- and low-frequency components of Radial Sway, decreasing the stability, and while additive noise is effective in decreasing RS in the baseline quiet condition and in the low-frequency component of sway, it was not effective in the high-frequency component.

This work was able to show the differential influences of both sensory input and cognitive demand on the dynamics of postural sway while offering insight into the possible utilization of additive sensory noise in environments where postural stability may suffer. These results were able to show the negative influence of cognitive load on postural sway, with higher cognitive load resulting in a decrease in stability. However, with the additive white noise, we were able to return stability back to baseline-like levels, implying that additive noise makes the body more stable even when performing a destabilizing task.

Cognitive load was predicted to increase sway variability regardless of the level of difficulty applied, while auditory noise was predicted to then reduce the variability back to normal levels, but this hypothesis was only partially true. The explanation behind our findings is unclear, but the implications are important. Noise interventions for improving balance may be relevant and effective regardless of the subsequent task at hand. If cognitive load causes an increase in sway variability, this additive auditory noise is able to reduce the sway back to normal settings seen prior to cognitive load. This may be critical in at risk populations, such as those with neurological disorders or ageing populations, to help aid in maintaining their motor performance during daily living.

Current theoretical explanation for this phenomenon of cognition and posture is lacking. However, for the influence of noise on postural sway, the theory of Stochastic Resonance (SR) is typically discussed. The theory of SR explains the amplification of information-carrying signals through the addition of broad-spectrum uncorrelated noise in a threshold-based system, such as and including the nervous system (Hangii 2002). A commonly held view of noise is that it obscures information within a signal and requires a filtering to increase the signal to noise ratio. However, evidence shows that noise can contribute to signal optimization in threshold-based systems specifically (Benzi et al. 1981; Hangii 2002). However, there is also the possibility that additive noise increases the attentional arousal during stimulation, which could lead to an increase in stability during standing. Cluff et al. (2010) showed that adding a cognitive task during standing leads to more automaticity in the balance process, which may improve stability due to the attention being shifted from the motor control of posture to the cognitive task at hand. This theory nevertheless has been disputed with work by Deviterne et al. (2005) in which they introduced a passive listening task of a single sustained auditory tone and found no effect of the stimuli on postural control.

As for the influence of cognitive load on the motor system, multiple ideas exist positing how there are a limited attentional resource available within the mind that must be allocated to the tasks that are being performed. When the more tasks performed, the less attentional resources each task is given, resulting in a reduction in performance of the tasks being performed simultaneously (Abou Khalil et al. 2020). Similarly, there are older psychological explanations that posit that there exists a predetermined limit and pathway of attention, and when multiple tasks are performed concurrently, a bottleneck occurs within the brain causing the interference effect we see in the behavioral data. Similarly, work by Abou Khalil et al. (2020) suggests that there is a preferential organization of the body for the optimization of cognitive performance, with walking having the most benefit for memory activity and sitting preferred for mental arithmetic. This work has led to an understanding of how the human sensory modalities influence cognitive attentional demands and how the change in attentional demands may interact with postural control dynamics. Past work has shown that any shift in conscious-controlled attention toward postural control increases the likelihood of disrupting coordination and stability (Masters and Maxwell 2008; Wulf et al. 2001). This disruption is commonly posited to be a consequence of reinvestment theory (Masters and Maxwell 2008; Masters 1992), which suggests that relatively automated motor processes can be disrupted if they are being consciously accessed, using task-relevant declarative knowledge (Masters 1992).

This theory also suggests that aging and neurological diseases cause an increase in reinvestment (Masters and Maxwell 2008; Schaefer et al. 2015). This was shown by Seidler et al. (2010), who found that physiological changes with aging and injury are due to loss in the gray and white matter within the CNS, resulting in differential-reorganized cortical activation (Ghai et al. 2017). Which led the authors to suggest that differential cortical activity within the higher neural centers can affect task prioritization, further allowing increased conscious attention while performing secondary motor or cognitive tasks (Talelli et al. 2008).

For healthy adults, standing is so over-practiced that it often seems automatic (Bernstein 1996; Fitts and Posner 1967). However, postural control is far from simple—it demands the coordination of nearly all major muscle groups. This coordination is thought to be achieved by forming a postural synergy (Bardy et al. 1999; Latash 1998), which operates under the guidance of multiple perceptual systems. Furthermore, postural control must adapt to suprapostural activities (Riccio and Stoffregen 1988) and be prepared to resist unexpected perturbations.

Considering these factors and the potential involvement of cortical processes, postural control is a complex, high-level neurocognitive feat.

Since postural control involves task-specific interactions between central processes, perception-action coupling, behavioral constraints, and environmental context, any of these factors can influence the composition and organization of postural synergies. Cognitive load, perceptual demands, suprapostural behaviors, and environmental conditions (such as the rigidity of the support surface) can all affect the emergence of task-specific postural synergies. This suggests a hypothesis regarding the relationship between cognition and postural control. Rather than cognitive load diminishing postural control by diverting attentional resources needed to manage postural synergies, it may instead constrain the organization of those synergies (see Jeka 1995; Mitra et al. 1998; Riley and Turvey 2002, for related discussions on perceptual constraints in coordination and synergies). This perspective can explain both increases and decreases in postural stability under cognitive load—something the resource allocation model struggles to do, particularly when explaining increases in stability during standing. This hypothesis aligns with Pellechia’s (2003) suggestion that concurrent postural and cognitive tasks may reflect higher-order coordination between the two activities (see Neumann 1996). The coordination of postural and cognitive activities could introduce constraints on the organization of postural synergies.

## Supporting information

Supplementary Statistics

## Data Availability

Datasets generated and analyzed during the current study are available from the corresponding author upon reasonable request.

## REFERENCES

Abou Khalil G, Doré-Mazars K, Senot P, Legrand A (2020) Is it better to sit down, stand up or walk when performing memory and arithmetic activities? Experimental Brain Research 238:2487–2496

Andersson G, Hagman J, Talianzadeh R, Svedberg A, Larsen HC (2002) Effect of cognitive load on postural control. Brain Res Bull 58:135–139

Bardy BG, Marin L, Stoffregen TA, Bootsma RJ (1999) Postural coordination modes considered as emergent phenomena. Journal of Experimental Psychology: Human Perception & Performance, 25:1284–1301

Benzi R, Sutera A, Vulpiani A (1981) The mechanism of stochastic resonance. J Phys A Math Gen 14:453–457

Bernstein N (1996) On dexterity and its development. In M. Latash & M. T. Turvey (Eds.), Dexterity and its development, 2–244. Mahwah, NK:Erlbaum

Blázquez MT, Anguiano M, Saavedra FA, Lallena AM, Carpena P (2010) Characterizing the human postural control system using detrended fluctuation analysis. J Comput Appl Mat 223:1478–1482

Boisgontier MP, Beets IA, Duysens J, Nieuwboer A, Krampe RT, Swinnen SP (2013) Agerelated differences in attentional cost associated with postural dual tasks: increased recruitment of generic cognitive resources in older adults. Neurosci Biobehav Rev 37(8):1824–1837

Carey S, Ross JM, Balasubramaniam R (2023) Auditory, tactile, and multimodal noise reduce balance variability. Experimental Brain Research 241(5):1241–1249

Carey S, Ross JM, Abney D, Balasubramaniam R (2024) Effects of auditory noise intensity and color on the dynamics of upright stance. Scientific Reports 14(1):10518

Centers for Disease Control and Prevention, National Center for Injury Prevention and Control. Web–based Injury Statistics Query and Reporting System (WISQARS) [online]

Cluff T, Gharib T, Balasubramaniam R (2010) Attentional influences on the performance of secondary physical tasks during posture control. Experimental Brain Research 203:647– 658. 10.1007/s00221-010-2274-7

Collins JJ, De Luca CJ (1994) Random walking during quiet standing. Phys Rev Lett 73:764– 767

Delignières D, Deschamps T, Legros A, Caillou N (2003) A methodological note on non-linear time series analysis: is Collins and De Luca (1993)’s open- and closed-loop model a statistical artifact? J Mot Behav 35:86–96

Deviterne D, Gauchard GC, Jamet M, Vançon G, Perrin PP (2005) Added cognitive load through rotary auditory stimulation can improve the quality of postural control in the elderly. Brain Res Bull 64:487–492

Dozza M, Horak FB, Chiari L (2007) Auditory biofeedback substitutes for loss of sensory information in maintaining stance. Experimental Brain Research 178:37–48

Duarte M, Freitas SM (2010) Revision of posturography based on force plate for balance evaluation. Brazilian Journal of physical therapy 14:183–192

Fitts PM, Posner MI (1967) Human Performance Belmont, CA: Brooks-Cole

Fraizer EV, Mitra S (2008) Methodological and interpretive issues in posture-cognition dualtasking in upright stance. Gait and Posture 27:271–279

Ghai S, Ghai I, Effenberg AO (2017) Effects of dual tasks and dual-task training on postural stability: a systematic review and meta-analysis. Clinical Interventions in Aging 12:557–577

Hänggi P (2002) Stochastic resonance in biology: how noise can enhance detection of weak signals and help improve biological information processing. ChemPhysChem 3:285–290

Hegeman J, Honegger F, Jupper M, Allum JHJ (2005) The balance control of bilateral peripheral vestibular loss subjects and its improvement with auditory prosthetic feedback. J Vestib Res 15:109–117

Honeycutt CF, Gottschall JS, Nichols TR (2009) Electromyographic responses from the hindlimb muscles of the decerebrate cat to horizontal support surface perturbations. J Neurophysiol 101(6):2751–2761

Hornbrook MC, Stevens VJ, Wingfield DJ, Hollis JF, Greenlick MR, Ory MG (1994) Preventing falls among community-dwelling older persons V results from a randomized trial. Gerontologist 34:16–23

Jacobs J, Horak F (2007) Cortical control of postural responses. J Neural Transm 114(10):1339– 1348

Jeka JJ (1995) Is servo-theory the language of human postural control? Ecological Psychology 7:321–327

Johansson JD, Lemon RN, Westling G (1994) Time varying enhancement of human cortical excitability mediated by cutaneous inputs during precision grip. Journal of Physiology 481:761–765

Kiemel T, Oie KS Jeka JJ (2005) Slow dynamics of postural sway are in the feedback loop. J Neurophysiol 95:1410–1418

Lafond D, Corriveau H, Hébert R, Prince F (2004) Intrasession reliability of center of pressure measures of postural steadiness in elderly people. Arch Phys Med Rehabilit 85:896–901

Lalonde R, Strazielle C (2007) Brain Regions and Genes Affecting Postural Control. Progress in Neurobiology 81:45–60

Latash ML (1998) Neurophysiological basis of movement. Urbana, UL: Human Kinetics

Lin D, Seol H, Nussbaum NA, Madigan ML (2008) Reliability of COP-based postural sway measures and age-related differences. Gait Posture 28:337–342

Masters RSW (1992) Knowledge, nerves and know-how: the role of explicit versus implicit knowledge in the breakdown of a complex motor skill under pressure. Br J Psychol 83(3), 343–358

Masters RSW, Maxwell J (2008) The theory of reinvestment. Int Rev Sport Exer Psychol 1(2):160–183

Maylor EA, Wing AM (1996) Age differences in postural stability are increased by additional cognitive demands. J Gerontol B-Psychol 51:143–54

Maylor EA, Allison S, Wing AM (2001) Effects of spatial and nonspatial cognitive activity on postural stability. Br J Psychol 92:319–338

Mitra S, Riley MA, Schmidt RC, Tuvey MT (1998) Vision and the level of synergies. In L.H. Harris & M. Jenkin (Eds.), Vision and action (pp. 314–331). New York: Cambridge University Press

Morasso P, Cherif A, Zenzeri J (2019) Quite standing: the single inverted pendulum model is not so bad after all. PLoS One 14(3):e0213870

Morton SM, Bastian AJ (2004) Cerebellar control of balance and locomotion. Neuroscientist 10:247–259

Neumann O (1996) Theories of attention. In O. Neumann & A. F. Sanders (Eds.), Handbook of perception and action, Vol 3: Attention (pp. 389–446). San Diego, CA: Academic Press

Pellecchia GL (2003) Postural sway increases with attentional demands of a concurrent cognitive task. Gait & Posture 18:29–34

Peng CK, Buldyrev SV, Havlin S, Simon M, Stanley HE, Golberger AL (1994) Mosaic organization of DNA nucleotides, Phys. Rev. E. 49:1685–1689

Priplata A, Niemi J, Salen M, Harry J, Lipsitz LA, Collins JJ (2002) Noise-enhanced balance control. Physics Review Letters 89:238101.1-238101.4

Priplata A, Niemei J, Harry J, Lipsitz L, Collins J (2003) Vibrating insoles and balance control in elderly people. Lance 362:1123–1124

Priplata AA, Patritti BL, Niemi JB, Hughes R, Gravele DC, Lipsitz LA, Veves A, Stein J, Bonato P, Collins J (2006) Noise-enhanced balance control in patients with diabetes and patients with stroke. Annual Neurology 59:4–12

Quant S, Adkin AL, Staines WR, Maki BE, McIlroy WE (2004) The effect of a concurrent cognitive task on cortical potentials evoked by unpredictable balance perturbations. BMC Neuroscience 5:18

Quant S, Maki BE, McIlroy WE (2005) The association between later cortical potentials and later phases of postural reactions evoked by perturbations to upright stance. Neuroscience Letters 381:269–74

Raftopoulos A (2005) Cognitive Penetrability of Perception: Attention, Action, Strategies, and Bottom-Up Constraints. New York, NY: Nova Publishers

Ramenzoni VC, Riley MA, Shockley K, Chiu CYP (2007) Postural responses to specific types of working memory tasks. Gait & Posture 25(3):368–373

Redfern MS, Jennings JR, Martin C, Furman JM (2001) Attention influences sensory integration for postural control in older adults. Gait & Posture 14:211–6

Riccio GE, Stoffregen TA (1988) Affordances as constraints on the control of stance. Human Movement Science 7:265–300

Riley MA, Turvey MT (2002) Variability and determinism in motor behavior. Journal of Motor Behavior 41:99–125

Ross JM, Balasubramaniam R (2015) Auditory white noise reduces postural fluctuations even in the absence of vision. Experimental Brain Research 233:2357–2363

Ross JM, Will OJ, McGann Z, Balasubramaniam R (2016a) Auditory white noise reduces age-related fluctuations in balance. Neuroscience Letters 630:216–221

Ross JM, Warlaumont AS, Abney DH, Rigoli LM, Balasubramaniam R (2016b) Influence of musical grove on postural sway. J Exp Psychol Hum Percept Perform 42(3):308–319

Seidler RD, Bernard JA, Burutolu TB, Fling BW, Gordon MT, Gwin JT, Kwak Y, Lipps DB (2010) Motor control and aging: links to age-related brain structural, functional, and biochemical effects. Neurosci Biobehav Rev 34(5):721–733

Schaefer S, Schellenbach M, Lindenberger U, Woollacott M (2015a) Walking in high-risk settings: do older adults still prioritize gait when distracted by a cognitive task? Experimental Brain Research 233(1):079–88

Stoffregen TA, Smart LJ, Bardy BG, Pagulayan RJ (1999) Postural stabilization of looking. Journal of Experimental Human Psychology 25:1641–58

Stoffregen TA, Pagulayan RJ, Bardy BG, Hettinger LJ (2000) Modulating postural control to facilitate visual performance. Human Movement Science 19:203–20

Talelli P, Ewas A, Waddingham W, Rothwell J, Ward N (2008) Neural correlates of age-related changes in cortical neurophysiology. Neuroimage 40(4):1772–1781

van den Heuvel MRC, Balasubramaniam R, Daffertshofer A, Longtin A, Beek PJ (2009) Delayed visual feedback reveals distinct time scales in balance control. Neuroscience Letters 452:37–41

Wikstrom EA, Tillman MD, Smith AN, Borsa PA (2005) A new force-plate technology measure of dynamic postural stability: the dynamic postural stability index. Journal of Athletic Training 40(4):305–309

Winter D, Patla A, Prince F, Ishac M, Gielo-Perczak K (1998) Stiffness control of balance in quiet standing. Journal of Neurophysiology 80:1211–1221

Wollacott M, Shumway-Cook A (2002) Attention and the control of posture and gait: a review of an emerging area of research. Gait & Posture 16:1–4

Wulf G, McNevin N, Shea CH (2001) The automaticity of complex motor skill learning as a function of attentional focus. Q J Exp Psychol A 54(4):1143–1154

Yeh TT, Boulet J, Cluff T, Balasubramaniam R (2010) Contributions of delayed visual feedback and cognitive task load to postural dynamics. Neuroscience Letters 481:173–177

Yeh TT, Cluf T, Balasubramaniam R (2014) Visual reliance for balance control in older adults persists when visual information is disrupted by artificial feedback delays. PLoS One 9:e91554

