## Supplementary Statistics for "Influence of Cognitive Demand and auditory noise on Postural Dynamics"

<http://www.rameshlab.com>

**Supplementary Information**

Radial Sway Post Hoc Comparisons:

| Contrasts | Estimate | SE | T-Ratio | P Value |
| --- | --- | --- | --- | --- |
| None Silent – Easy Silent | -0.1444 | -0.199 | -0.726 | 0.9969 |
| None Silent – Hard Silent | -0.7532 | -0.199 | -3.786 | 0.0023 |
| None Silent – None Noise | 1.2728 | 0.203 | 6.281 | <.0001 |
| None Silent – Easy Noise | 0.2979 | 0.200 | 1.493 | 0.6690 |
| None Silent – Hard Noise | 0.0613 | 0.200 | 0.307 | 0.9996 |
| Easy Silent – Hard Silent | -0.6087 | 0.195 | -3.144 | 0.0234 |
| Easy Silent – None Noise | 1.4172 | 0.200 | 7.095 | <.0001 |
| Easy Silent – Easy Noise | 0.4423 | 0.196 | 2.255 | 0.2142 |
| Easy Silent – Hard Noise | 0.2057 | 0.196 | 1.048 | 0.9013 |
| Hard Silent – None Noise | 2.0259 | 0.200 | 10.133 | <.0001 |
| Hard Silent – Easy Noise | 1.0510 | 0.196 | 5.356 | <.0001 |
| Hard Silent – Hard Noise | 0.8145 | 0.196 | 4.151 | 0.0005 |
| None Noise – Easy Noise | -0.9749 | 0.201 | -4.862 | <0.0001 |
| None Noise – Hard Noise | -1.2115 | 0.201 | -6.035 | <0.0001 |
| Easy Noise – Hard Noise | -0.2366 | 0.197 | -1.201 | 0.8363 |

High-Frequency RS Post Hoc Comparison:

| Contrasts | Estimate | SE | T-Ratio | P Value |
| --- | --- | --- | --- | --- |
| None Silent – Easy Silent | 0.0182 | 0.0755 | 0.242 | 0.9999 |
| None Silent – Hard Silent | -.3232 | 0.0760 | -4.251 | 0.0004 |
| None Silent – None Noise | 0.3472 | 0.0767 | 4.527 | 0.0001 |
| None Silent – Easy Noise | 0.1664 | 0.0758 | 2.195 | 0.2415 |
| None Silent – Hard Noise | -0.1335 | 0.0758 | -1.762 | 0.4909 |
| Easy Silent – Hard Silent | -0.3414 | 0.0746 | -4.578 | 0.0001 |
| Easy Silent – None Noise | 0.3290 | 0.0755 | 4.357 | 0.0002 |
| Easy Silent – Easy Noise | 0.1482 | 0.0743 | 1.994 | 0.3467 |
| Easy Silent – Hard Noise | -0.1518 | 0.0743 | -2.041 | 0.3200 |
| Hard Silent – None Noise | 0.6704 | 0.0760 | 8.816 | <.0001 |
| Hard Silent – Easy Noise | 0.4896 | 0.0749 | 6,537 | <.0001 |
| Hard Silent – Hard Noise | 0.1897 | 0.0747 | 2.540 | 0.1142 |
| None Noise – Easy Noise | -.1808 | 0.0758 | -2.383 | 0.1636 |
| None Noise – Hard Noise | -0.4807 | 0.0758 | -6.344 | <.0001 |
| Easy Noise – Hard Noise | -0.3000 | 0.0747 | -4.016 | 0.0009 |

Low-Frequency RS Post Hoc Comparisons:

| Contrasts | Estimate | SE | T-Ratio | P Value |
| --- | --- | --- | --- | --- |
| None Silent – Easy Silent | 0.0867 | 0.181 | 0.479 | 0.9969 |
| None Silent – Hard Silent | -0.5236 | 0.180 | -2.903 | 0.0441 |
| None Silent – None Noise | 1.0728 | 0.184 | 5.826 | <.0001 |
| None Silent – Easy Noise | 0.5048 | 0.182 | 2.774 | 0.0629 |
| None Silent – Hard Noise | 0.3494 | 0.181 | 1.933 | 0.3827 |
| Easy Silent – Hard Silent | -0.6103 | 0.178 | -3.437 | 0.0082 |
| Easy Silent – None Noise | 0.9860 | 0.182 | 5.418 | <.0001 |
| Easy Silent – Easy Noise | 0.4180 | 0.179 | 2.335 | 0.1816 |
| Easy Silent – Hard Noise | 0.2627 | 0.178 | 1.477 | 0.6791 |
| Hard Silent – None Noise | 1.5963 | 0.181 | 8.814 | <.0001 |
| Hard Silent – Easy Noise | 1.0283 | 0.178 | 5.764 | <.0001 |
| Hard Silent – Hard Noise | 0.8730 | 0.177 | 4.928 | <.0001 |
| None Noise – Easy Noise | -0.5680 | 0.183 | -3.107 | 0.0241 |
| None Noise – Hard Noise | -0.7234 | 0.182 | -3.983 | 0.0011 |
| Easy Noise – Hard Noise | -0.1553 | 0.179 | -0.869 | 0.9536 |

DFA Post Hoc Comparisons:

| Contrasts | Estimate | SE | T-Ratio | P Value |
| --- | --- | --- | --- | --- |
| None Silent – Easy Silent | 0.01948 | 0.0115 | 1.695 | 0.5356 |
| None Silent – Hard Silent | 0.03108 | 0.0116 | 2.686 | 0.0791 |
| None Silent – None Noise | 0.09826 | 0.0117 | 8.363 | <.0001 |
| None Silent – Easy Noise | 0.05501 | 0.0116 | 4.738 | <.0001 |
| None Silent – Hard Noise | 0.05669 | 0.0116 | 4.899 | <.0001 |
| Easy Silent – Hard Silent | 0.01160 | 0.0114 | 1.016 | 0.9126 |
| Easy Silent – None Noise | 0.07878 | 0.0116 | 6.766 | <.0001 |
| Easy Silent – Easy Noise | 0.03553 | 0.0115 | 3.102 | 0.0243 |
| Easy Silent – Hard Noise | 0.03721 | 0.0114 | 3.259 | 0.0148 |
| Hard Silent – None Noise | 0.06718 | 0.0117 | 5.735 | <.0001 |
| Hard Silent – Easy Noise | 0.02393 | 0.0115 | 2.076 | 0.3012 |
| Hard Silent – Hard Noise | 0.02561 | 0.0115 | 2.232 | 0.2245 |
| None Noise – Easy Noise | -0.04325 | 0.0118 | -3.681 | 0.0034 |
| None Noise – Hard Noise | -0.04157 | 0.0117 | -3.546 | 0.0055 |
| Easy Noise – Hard Noise | 0.00168 | 0.0115 | 0.146 | 1.0000 |
